# Short latency stretch reflexes depend on the balance of activity in agonist and antagonist muscles during ballistic elbow movements

**DOI:** 10.1101/2021.12.06.471376

**Authors:** Zoe Villamar, Daniel Ludvig, Eric J Perreault

## Abstract

The spinal stretch reflex is a fundamental building block of motor function, with a sensitivity that varies continuously during movement and when changing between movement and posture. Many have investigated task-dependent reflex sensitivity, but few have provided simple, quantitative analyses of the relationship between the volitional control and stretch reflex sensitivity throughout tasks that require coordinated activity of several muscles. Here we develop such an analysis and use it to test the hypothesis that modulation of reflex sensitivity during movement can be explained by the balance of activity within agonist and antagonist muscles better than by activity only in the muscle homonymous with the reflex. Subjects completed hundreds of flexion and extension movements as small, pseudo-random perturbations of elbow angle were applied to obtain estimates of stretch reflex amplitude throughout the movement. A subset of subjects performed a postural control task at muscle activities matched to those during movement. We found that reflex modulation during movement can be described by background activity in antagonist muscles about the elbow much better than by activity only in the muscle homonymous to the reflex (p<0.001). Agonist muscle activity enhanced reflex sensitivity whereas antagonist activity suppressed it. Surprisingly, the magnitude of these effects was similar, suggesting a balance of control between agonists and antagonists very different from the dominance of sensitivity to homonymous activity during posture. This balance is due to a large decrease in sensitivity to homonymous muscle activity during movement rather than substantial changes in the influence of antagonistic muscle activity.

**New and Noteworthy:** This study examined the sensitivity of the stretch reflexes elicited in elbow muscles to the background activity in these same muscles during movement and postural tasks. We found a heightened reciprocal control of reflex sensitivity during movement that was not present during maintenance of posture. These results help explain previous discrepancies in reflex sensitivity measured during movement and posture and provide a simple model for assessing their contributions to muscle activity in both tasks.

## Introduction

The spinal stretch reflex is a fundamental building block of motor function, modulating its sensitivity across tasks to augment volitional control. The sensitivity of stretch reflexes varies continuously during movement (Stein Richard B. et al., 1988; Zehr et al., 1997), differs between posture and movement (Bawa et al., 1999; Brooke et al., 1997), and can vary with fusimotor set (Prochazka et al., 2012). While there have been many demonstrations of reflex modulation and investigations into the underlying mechanisms, there have been few attempts to provide simple, quantitative descriptions of the relationship between the volitional control and stretch reflex sensitivity throughout tasks that require the coordinated activity of several muscles. Here we develop such a description, providing a framework for understanding the continuous modulation of stretch reflexes in unimpaired subjects and a baseline for quantifying changes due to injury, pathology, or aging.

It is well documented that the spinal stretch reflex exhibits gain-scaling, whereby its sensitivity to external perturbations of posture is proportional to the background activity in the stretched muscle (Bawa, et al., 1999; Cathers et al., 2004; Matthews, 1986). Gain-scaling in homonymous stretch reflexes is an important example of coordination between volitional and reflexive motor control, presumably regulated through *α-γ* co-activation (Burke et al., 1978; Burke et al., 1978; Hagbarth et al., 1969; Vallbo, 1970), activation of *β* motoneurons (Bessou et al., 1963; Emonet-Denand et al., 1975), or increased recruitment of progressively larger motor units (Henneman et al., 1965; Henneman et al., 1965). Gain-scaling can be used to characterize activation-dependent sensitivity of the stretch reflex within individual postural tasks (Matthews, 1986; Pruszynski et al., 2009; Stein R. B. et al., 1995), but often fails when assessed across similar postural tasks requiring different patterns of volitional muscle coordination (Sohn et al., 2017) or during movement (Nakazawa et al., 1997; Soechting et al., 1981; Wallace et al., 1998). Recently, Dimitriou (2014) used microneurography to demonstrate that muscle spindle activity recorded during movement cannot be explained by simple gain-scaling, but rather represents the balance of activity within agonist and antagonist muscles crossing a joint. If this balance extends to the level of whole-muscle stretch reflexes, it could be useful means to describe task-dependent modulation in the spinal stretch reflex.

In addition to being modulated continuously, reflex sensitivity decreases during movement relative to the sensitivity observed during posture. In an elegant series of experiments, Brooke and colleagues demonstrated that at least some of this decrease arises from presynaptic inhibition from sensory afferents (Brooke et al., 1995; Brooke et al., 1995). However, most movements require the coordinated activation of agonists and antagonists that is not always required for the laboratory tasks often used to study postural stretch reflexes (Lacquaniti et al., 1982). Hence, the balance of activity that regulates muscle spindle sensitivity may also contribute to the commonly observed reduction in stretch reflex sensitivity during movement along with modulation of that sensitivity throughout movement. Such a mechanism could amplify the effects of afferent-mediated presynaptic inhibition (Macefield et al., 2018). A challenge in testing this possibility has been obtaining continuous estimates of reflex sensitivity throughout the course of movement, as is helpful for a rich enough dataset to explore the factors contributing to reflex modulation.

The objective of this study was to test the hypothesis that the modulation of stretch reflex sensitivity during movements could be explained by the balance of activity within the relevant agonist and antagonist muscles. This was accomplished by comparing a model that considered the effects of agonist and antagonist background activity on reflex amplitudes to one that considered only simple gain-scaling from the homonymous muscle. All assessments were made within muscles spanning the human elbow as subjects made repeated elbow flexion and extension movements. We used an innovative experimental approach that allowed us to obtain continuous estimates of stretch reflex amplitude throughout the movement. This was done by applying continuous pseudo-random perturbations of elbow angle throughout the movement, and collecting data from approximately 500 movements in each direction so averaged estimates of background muscle activity and reflex amplitude could be obtained throughout the movement. We also ran a control experiment on a subset of subjects to determine in the model describing reflex modulation during movement could also describe the activity-dependent modulation often reported during postural control, when assessed at similar levels of background activity in both tasks. Our results demonstrate that the balance of activity contributing to muscle spindle sensitivity during movement can be observed at the level of whole-muscle stretch reflexes, but that this balance is not sufficient to explain differences between reflex activity in posture and movement control.

## Methods

### Participants

Eighteen individuals (11 females) between the ages 22 and 32 volunteered to participate in this study. Fourteen of these individuals (7 females) participated in the movement experiment only. Four additional individuals participated in a postural control experiment in addition to the movement experiment. All participants were neurologically intact and had no musculoskeletal impairments in the upper extremity. All participants were right-arm dominant. Ethical approval for the study was received from the Northwestern University Institutional Review Board. Written informed consent was obtained prior to testing.

### Experimental Set-Up

A rotary motor aligned with the elbow was used to apply perturbations necessary to elicit stretch reflexes as well as create a virtual environment that allowed subjects to perform consistent natural reaching movements. Participants were seated in a height-adjustable chair (Biodex, Shirley, NY) and were secured to the chair with straps across the torso. The participant’s right arm was placed in a posture 70 degrees abducted in the frontal plane, 45 degrees of shoulder flexion in the transverse plane, and elbow flexion of 115 degrees. In this posture, the subject’s arm was attached to the electric rotary motor (BSM90N-3150AF, Baldor, Fort Smith, AR), via a rigid fiberglass cast. The fiberglass forearm cast extended from the palm to a point just distal of the elbow (**Figure 1**). The center of rotation of the elbow was aligned to the center of rotation of the motor. To prevent the motor pushing the elbow beyond the voluntary range of motion, electrical and mechanical safety stops were placed within each participant’s voluntary range of motion.

**Figure 1.**
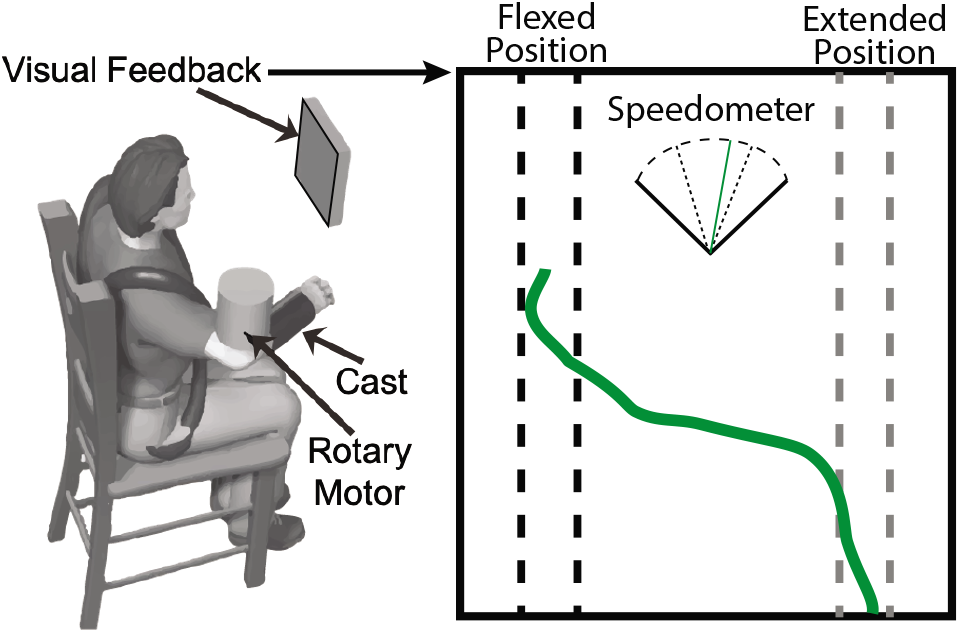
Experimental Set-Up and Visual Feedback. Subjects were seated with their elbow attached to a rotary motor via a fiberglass cast. For the movement task, the subject’s position was represented by the green line. The current position was at the top-most point of the line, followed by 3 seconds of position history. The target was given by the two channels on the left (flexed position) and right side (extended) of the screen, with the black representing the current target. The speedometer between the channels provided speed feedback of the movement, with a green line between the dashed bounds representing the past movement matching the target speed.

Surface EMG was recorded from two elbow flexors: brachioradialis (BRD) and biceps brachii (BIC); and two extensors: triceps long head (TRILO) and triceps lateral head (TRILAT). Standard skin preparation methods (Tankisi et al., 2020) were applied prior to the application of bipolar electrodes (Noraxon 272, Scottsdale, AZ). EMG measurements were amplified (AMT8, Bortec, Calgary, AB) as needed to maximize the range of the data acquisition system. EMG data was filtered at 500 Hz with a 5-pole Bessel filter and sampled at 2.5 kHz (PCI-DAS1602/16, Measurement Computing, Norton, MA, USA). Elbow angle was measured with an encoder built into the rotary motor recorded simultaneously with the EMG using a 24-bit quadrature encoder card (PCI-QUAD04, Measurement Computing, Norton, MA, USA). Data acquisition, rendering of the virtual environment, and control of the rotary motor were executed using xPC target (The Mathworks Inc., Natick, MA).

### Protocol: Movement Task

The first experiment was designed to determine how reflex sensitivity varies with the background activity of various muscles during movement. This experiment involved repeated ballistic elbow flexion and extension movements as the participant interacted with a virtual environment. The virtual environment consisted of an inertia of 1 kgm^2^, allowing us to simulate natural reaching with a modest load. A flexion torque bias equivalent to 5% of the participant’s maximum voluntary isometric torque was used to help maintain a consistent, non-zero level of flexion activity during the postural phase of each task. Subjects moved ballistically between 105 and 125 degrees of elbow extension, a range of 0.35 rad; full extension was defined as 180 degrees. The target angle switched after 1 second of being within 0.1 rad of the target posture (**Figure 1**). Visual feedback of position and speed was provided to help subjects perform consistent movements. Rotational perturbations were imposed during the movement to stretch the muscles crossing the elbow and elicit stretch reflexes within them. We used continuous stochastic perturbations, similar to several previous studies (Alibiglou et al., 2008; Kearney et al., 1997; Krutky et al., 2012; Ludvig et al., 2017). The perturbations we used were pseudorandom binary sequences with an amplitude 0.03 rad, a switching time 0.150 seconds, and a velocity 1.5 rad/second. These short duration perturbations accentuate short latency stretch reflexes (Lewis et al., 2005), which are the focus of this work. Twenty trials, each lasting 185 seconds, were conducted; each trial contained approximately 27 voluntary movements in the flexion and extension directions. Prior to the start of the experiments, isometric maximum voluntary contractions (MVC) were performed in elbow flexion and extension to later normalize to EMG and compare across participants. One participant of the movement experiment was removed from the analysis due to poor EMG recordings.

### Protocol: Postural Control Task

A second set of experiments was completed to compare reflex sensitivity between postural control and movement conditions. We designed a postural control task that would elicit stretch reflexes over a comparable range of background muscle activations as was encountered in the movement task described above. The motor was again configured with the same compliant environment used in the movement tasks, though subjects were now instructed to maintain a constant elbow posture. Several bias torques were used to control agonist muscle activity. Subjects were given real-time visual feedback of elbow angle and EMG in the antagonist muscle, allowing us to also specify different levels of co-contraction. For trials with a flexion bias, the EMG signal for feedback was from the TRILAT. For trials with an extension bias, the EMG signal used for feedback was from the BRD. Bias torques ranged from 15% MVC in flexion to 10% MVC in extension in increments of 5% MVC. For each level of bias torque, antagonist EMG target was either 0% MVC or 5% MVC. Each postural control trial lasted 65 seconds and was repeated twice.

### Data Analysis

Reflex activity was quantified at each time point within the average movement profile. Flexion and extension movements were processed independently. EMG data were notch filtered at 60 Hz then demeaned and rectified. EMG recordings for each muscle were normalized by the corresponding EMG recorded in the MVC trials (Besomi et al., 2020). The angle (**Figure 2A**) and EMG data (**Figures 2B, 2C**) were segmented into 8-second segments to create an ensemble of up to 350 segments per movement direction. To align these segments, we performed a correlation between each movement profile and a ramp signal representing a ballistic movement of 0.4 radians within 0.4 seconds. We subsequently aligned each segment using the time-shift that produced the peak correlation. Identical shifts were used for the EMG segments as were used for the corresponding movement segments.

**Figure 2.**
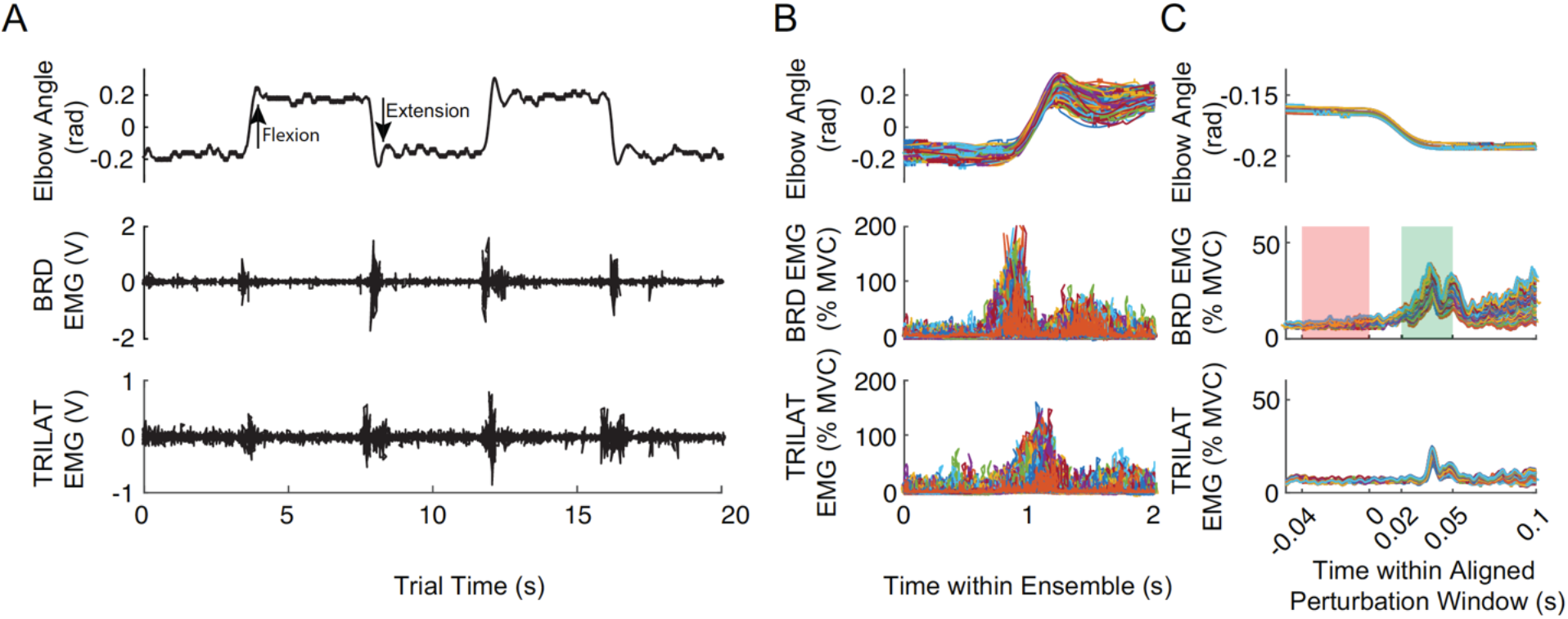
Metrics obtained to quantify muscle background and stretch reflex. **A,** Raw elbow angle recording as typical subject flexed and extended their elbow while small position perturbations were applied, along with the corresponding raw surface EMG signal from the BRD and the TRILAT muscles. **B**, ensemble of the segmented and aligned data for the elbow angle (top), BRD (middle), TRILAT (bottom) for a flexion movement, with the different colors representing each movement. **C**, a sample window from the ensemble in which the extension perturbations (top) and evoked responses for BRD (middle) and TRILAT (bottom) were aligned to the small perturbations superimposed on the larger voluntary movement. Shaded areas indicate where the background activity (red) and reflex activity (green) were quantified. Only muscle responses to imposed lengthening were considered.

To quantify reflexes along the ensemble, perturbations and their evoked responses were identified. Only responses evoked by perturbations that caused muscle lengthening were considered. Responses within a 200-ms window, centered at each timepoint of the ensemble, were averaged to estimate the reflex responses at each point along the voluntary movement trajectory (**Figure 2**). For each response, background EMG was defined as the average activity from 0-40 ms prior to perturbation onset. Short latency stretch reflexes were quantified by the average EMG 20 – 50 ms following the perturbation (Shemmell et al., 2010). The background activity was subtracted from the stretch reflex activity to get a relative measure of the stretch reflex activity.

We were interested in the time-course of stretch reflex activity and how it was influenced by background muscle activity throughout the duration of each movement. To simplify our subsequent analysis and data presentation, we compensated for delays in sensorimotor processing by shifting the time windows used for averaging background muscle activity and stretch reflexes so that both centered about the perturbation onset linking these quantities. The averaged background activity was shifted forward by 20 ms and the averaged reflex activity was shifted backward by 35 ms. This same process was used in the postural control task for consistency of processing, though it was not critical since those measures were made under steady-state conditions.

### Statistics

Our first goal was to determine the contributions from agonist and antagonist muscles to the amplitude of the short latency stretch reflex during movement. Our hypothesis was that stretch reflex modulation in a given target muscle could be predicted by activity in the target and its antagonist muscles better than when considering only the activity in the target and synergist. We fitted a linear mixed effects model for each muscle with the short latency stretch reflex activity as the dependent variable, the background activity in all four muscles as continuous factors, movement direction as a fixed factor, and subject as a random factor. We tested our hypothesis using a likelihood ratio to evaluate if inclusion of the background activity in the antagonist muscles allowed for significantly better prediction of stretch reflex activity during movement compared to the model without the antagonist muscles. We evaluated the change in variance accounted for after removing the contributions from the random effects for each subject. These analyses were performed on data decimated to 12.5 Hz to reduce correlation between sequential values in the time series and avoid inflated p-values (Neter et al., 1996), we decimated the data to 12.5 Hz. In practice, this had little effect since correlations were already considered by the structure of our linear mixed effects model. The 12.5 Hz we chose is similar to the sampling rate used in previous studies using spindle recordings rather than whole muscle EMG (Dimitriou, 2014).

We used a similar analysis to estimate the gain between background activity and the reflex amplitude in the target to evaluate how reflexes depended on activity in the agonist and antagonist muscles. Since the EMGs in synergistic muscles are highly correlated, we included only the background activity from the target muscle and one antagonist. For the models of flexor reflexes, the TRILAT was chosen as the antagonist. For models of extensor reflexes, the BRD was chosen as the antagonist. The TRILAT was chosen because it is a uniarticular muscle and the reflex traces typically were less noisy than for the TRILO. The BRD was chosen because it is also a uniarticular muscle and the electrode was located on the forearm, away from the belly of the other muscles in the upper arm, reducing the likelihood of crosstalk. The significance of the estimated gains between from target and antagonist muscles to the stretch reflex were determined using a Wald t-statistic with degrees of freedom estimated by a Satterthwaite approximation (Luke, 2017).

In addition to our primary hypothesis related to movement, we ran a control experiment to evaluate if the influence of background activity on the reflexes elicited during movement was similar to that observed during the maintenance of posture. This was accomplished using data from the four subjects who completed the movement and posture protocols. Two statistical tests were conducted. First, we tested if there was a difference in reflex amplitude at matched levels of background activity using linear mixed effects model, with either the background (to determine if levels of background activity were matched) or reflex (to determine task differences in reflex amplitude) as the dependent variable and the task (posture vs movement) as a fixed factor. We evaluated if there was a significant fixed effect of task using a Wald t-statistic with degrees of freedom estimated using a Satterthwaite approximation. Second, we compared the gains between the background and reflex EMGs. This was done using the same linear mixed-effects models as for our primary hypothesis, but instead of having movement direction (flexion vs. extension) as a fixed factor, we had task (posture vs. movement) as a fixed factor, since we found that the effect of movement direction was modest when testing our primary hypothesis. The differences between posture and movement were again assessed using a Wald t-statistic. All values are reported as the estimate ± the standard error.

## Results

During movement, there was modulation in the reflex activity of each muscle not observed in its background activity. Typical tri-phasic bursts were observed in the background activity, with the agonist muscle showing a burst of activity, followed by a braking burst in the antagonist, and a final corrective burst in the agonist muscle (**Figure 3**). These patterns were observed for flexion and extension movements. Reflex activity in the agonist muscle for each movement direction increased immediately prior to movement, as shown in the BRD for flexion movements and the TRILAT for extension movements (**Figure 3D,E**). Stretch reflexes in these muscles then dropped precipitously during movement before returning to the steady-state level measured during the hold phase between successive flexion and extension movements. These changes in reflex activity could not be attributed solely to the modulation of background activity in either the BRD or TRILAT (**Figure 3B,C**), as neither showed a similar cessation of activity. When looking at a specific muscle’s reflex activity, we refer to it as the target muscle. Across all subjects and both movement directions, the target muscle’s background activity was only moderately predictive of its reflex activity, accounting for 72% of the reflex activity variance in BRD, 61% in the BIC, 69% in TRILO and 72% in TRILAT.

**Figure 3.**
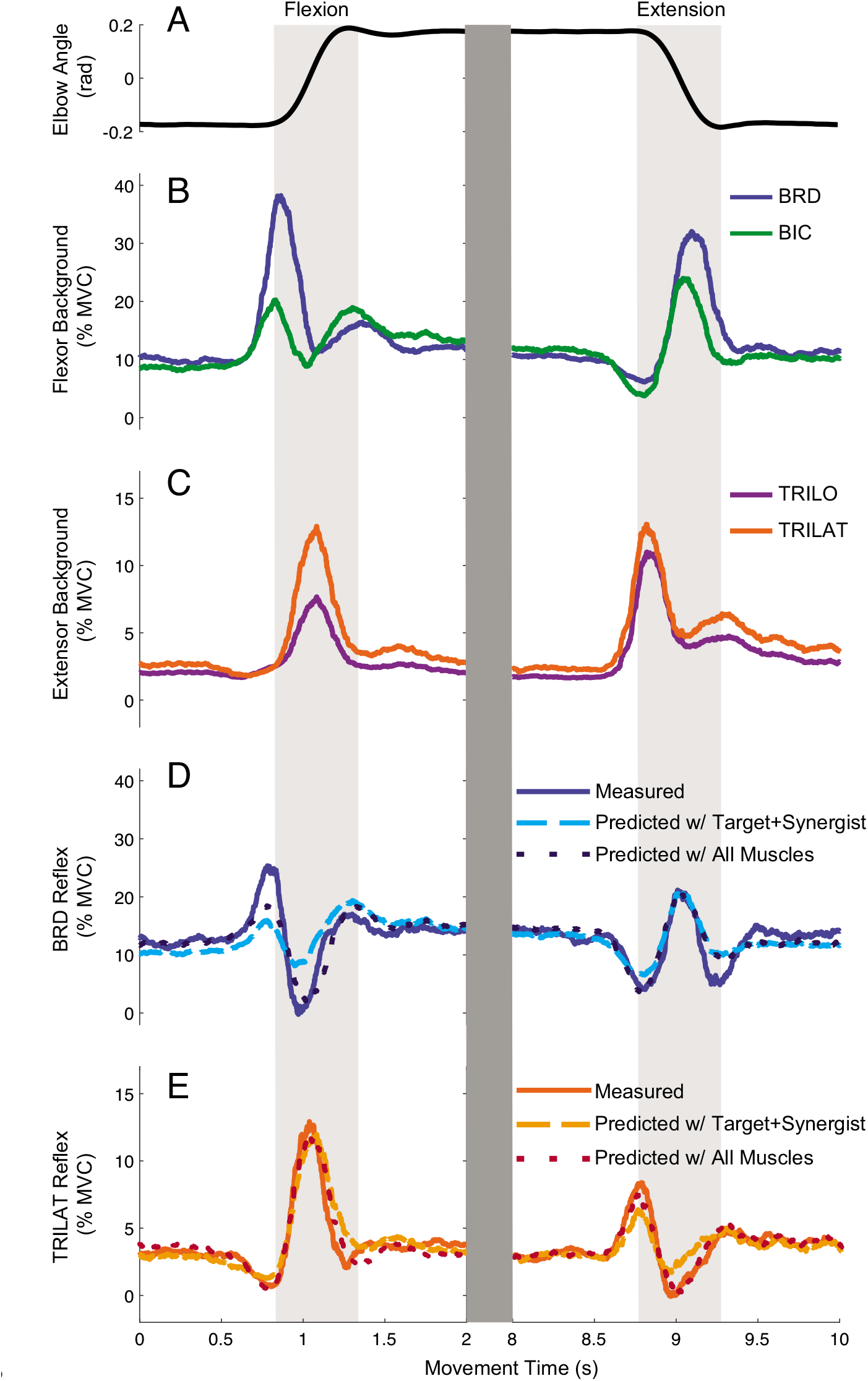
Representative data for an individual subject during flexion and extension movement. **A**) Trace of elbow position during flexion and extension movement; **B**) flexor muscle background activity; **C**) extensor muscle background activity; **D**) BRD reflex activity; **E**) TRILAT reflex activity.

During movement, stretch reflexes were sensitive to the background activity of both the agonist and antagonist muscles acting about the elbow. **Figure 3D-E** shows the reflex activity for a representative subject that was predicted using the background activity of the target muscle and its synergist, as well as one that was predicted using the background activity of the target, synergist, and antagonist muscles. For this representative subject, reflex prediction improved with the inclusion of the antagonist muscle backgrounds in the model for both the BRD muscle (R^2^ increased from 0.50 to 0.76) and TRILAT muscle (R^2^ increased from 0.80 to 0.91). With the inclusion of the antagonist muscle backgrounds, there was a very good prediction of the reflex, particularly in the TRILAT. For all muscles across all subjects, inclusion of the background activity of the antagonist muscles significantly improved the ability to predict reflex activity, accounting for approximately 25% more of the variance across all tested muscles (**Table 1**).

**Table 1:** R^2^ and Likelihood Ratio Statistics for the Models Predicting Reflexes

| **Muscle** | **$R^2$**<br>**(target+**<br>**synergist)** | **$R^2$ (2 synergists + 2**<br>**antagonists)** | **Likelihood**<br>**Ratio** | **Degrees**<br>**of**<br>**Freedom**<br>**(DoF)** | **p-**<br>**value** |
| --- | --- | --- | --- | --- | --- |
| **BRD** | 0.72 | 0.89 | 518 | 38 | <0.001 |
| **BIC** | 0.61 | 0.83 | 457 | 38 | <0.001 |
| **TRILO** | 0.69 | 0.83 | 437 | 38 | <0.001 |
| **TRILAT** | 0.72 | 0.89 | 656 | 38 | <0.001 |

Increased background activity in the antagonist muscle was associated with a reduction of reflex activity in the target muscle. Due to the background activity of the two flexor muscles being correlated (r = 0.50, p<0.001), as well as the background activity of the two extensor muscles being correlated (r = 0.71, p<0.001), it was not possible to obtain accurate estimates of the sensitivity of the reflex to any particular muscle. We therefore simplified our analysis to include only the BRD and TRILAT. Figure 4 shows how the reflex amplitude in each of these muscles depended on the background activities in both, a dependency we quantify by the gains between the background activities and reflex amplitude. These values were estimated using the linear mixed effects models described above. For the BRD reflex, its background had a positive gain of 0.54 ± 0.06 (t_14_ = 9.0, p < 0.001) while the TRILAT background activity, its antagonist, had a gain of −0.38 ± 0.09 (t_13_ = −4.1, p = 0.001) (**Figure 4A**). For the TRILAT reflex, its background had a positive gain of 0.49 ± 0.12 (t_14_ = 4.2, p < 0.001) while the BRD, its antagonist, had a gain of −0.32 ± 0.07 (t_9_ = −4.7, p < 0.001) (**Figure 4B**). Interestingly, these data point to a clear suppressive effect of the antagonist of nearly the same magnitude as the excitatory influence of the target muscle background activity.

**FIGURE 4.**
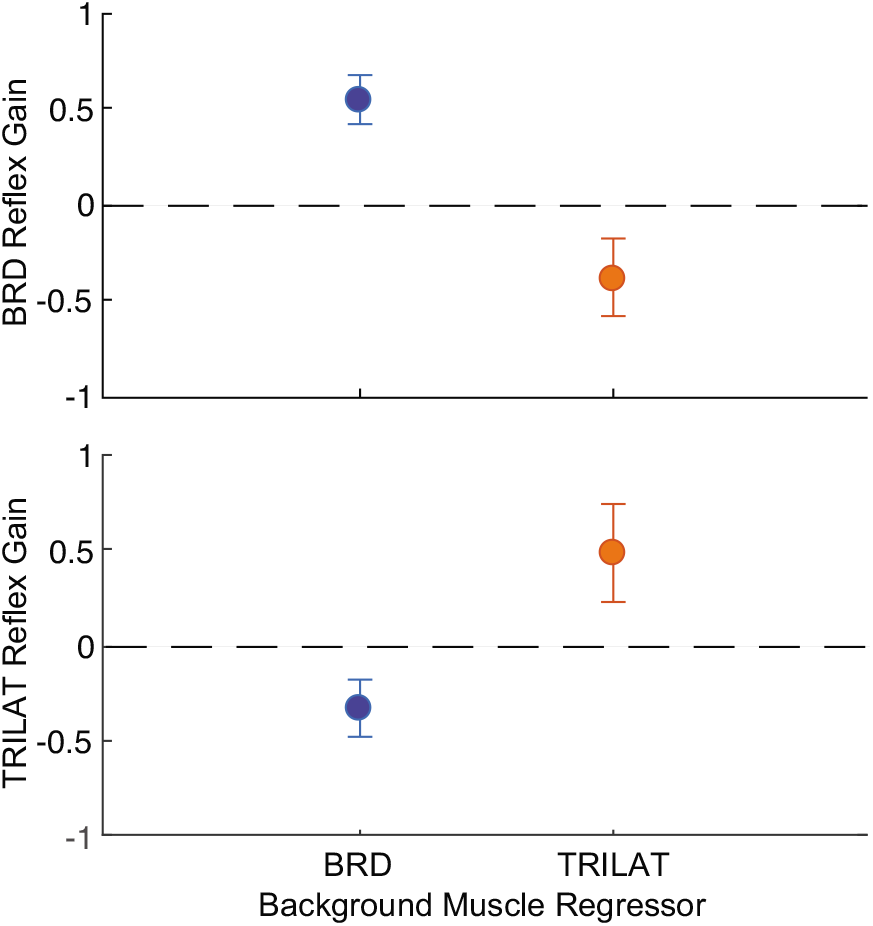
Gains of target muscle background and antagonist muscle background for **A**, a flexor and **B**, extensor during movement, for all participants who completed the movement trials (N = 17). Error bars are 95% confidence intervals. For the target muscle background for each reflex model, the gain is positive. For the antagonist muscle background for each reflex model, the gain is negative.

The above results are averaged across movement direction, which we found had only a modest effect on the estimated gains. The largest effect was found for the influence of the BRD background activity on its own reflex during flexion (0.72 ± 0.09) compared to extension (0.37 ± 0.09, t_9.1_ = 2.75, p = 0.02). The influence of the TRILAT background on the BRD reflex varied less with movement direction (Flex: −0.32 ± 0.10; Ext: −0.44 ± 0.10, t_8.1_ = 1.82, p = 0.11). The TRILAT reflex had quite modest differences in sensitivity to its background activity (Flex: 0.46 ± 0.12; Ext: 0.52 ± 0.13; t_9.3_ = −0.77, p = 0.46) or that of the BRD (Flex: −0.24 ± 0.05; Ext: −0.40 ± 0.10; t_8.2_ = 2.16, p = 0.06) during extension compared to flexion (**Figure 5**). For these reasons, movement direction was omitted as an independent variable in our control experiment used to assess differences between posture and movement.

**Figure 5.**
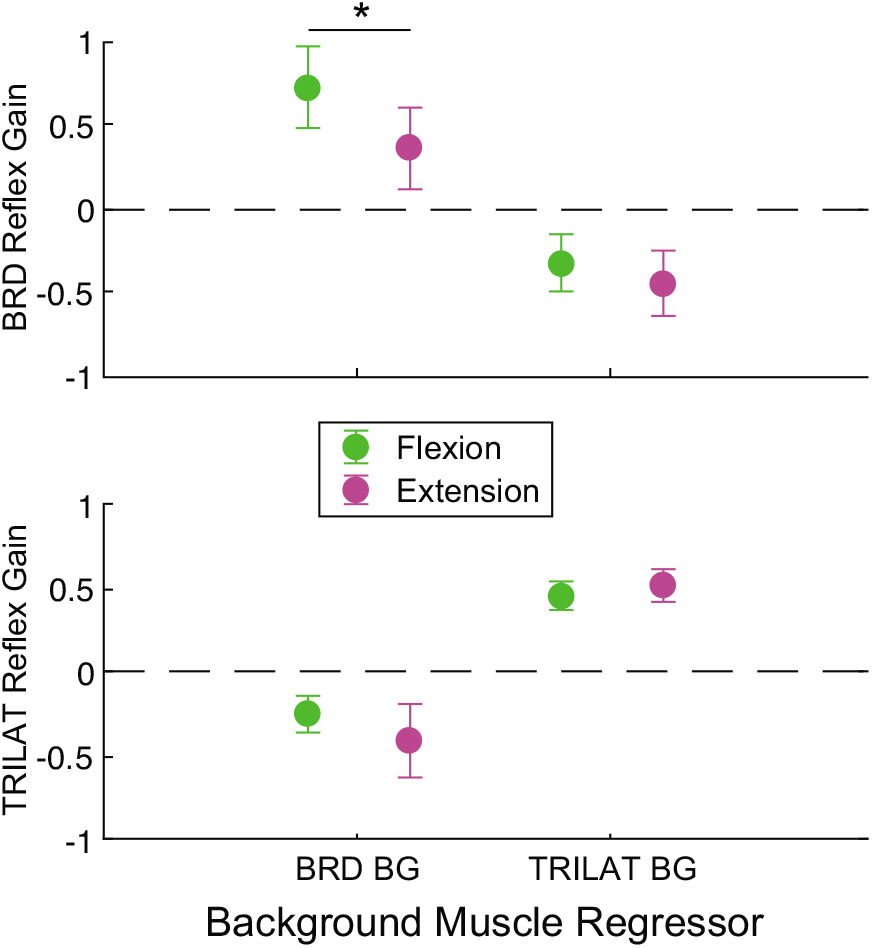
The effect of movement direction (flexion vs extension) on the gains of target muscle background and antagonist muscle background for a flexor and extensor reflex response, for all participants who completed the movement trials (N = 17). Error bars are 95% confidence intervals. * signifies p < 0.05.

The reflexes elicited during movement were smaller than those elicited during the maintenance of posture, as noted in several previous studies (Bawa, et al., 1999; MacKay et al., 1983). This was assessed by evaluating the task effect on the reflex amplitude at matched levels of background activity (**Figure 6**). During movement, the BRD reflex amplitude decreased by 5.11 ± 1.25 %MVC (t_3.4_ = −4.08, p = 0.02) and the TRILAT reflex amplitude decreased by 9.45 ± 1.04 %MVC (t_5.4_ = −9.01, p <0.001). These represent drops of 39% and 76%, respectively. These changes in reflex amplitude occurred even though the background activity was matched across tasks (BRD: Δ_Background_ = 0.39 ± 0.47 %MVC, t_45_ = 0.83, p = 0.41; TRILAT: Δ_Background_ = 0.26 ± 0.56 %MVC, t_63_ = 0.47, p = 0.64).

**Figure 6.**
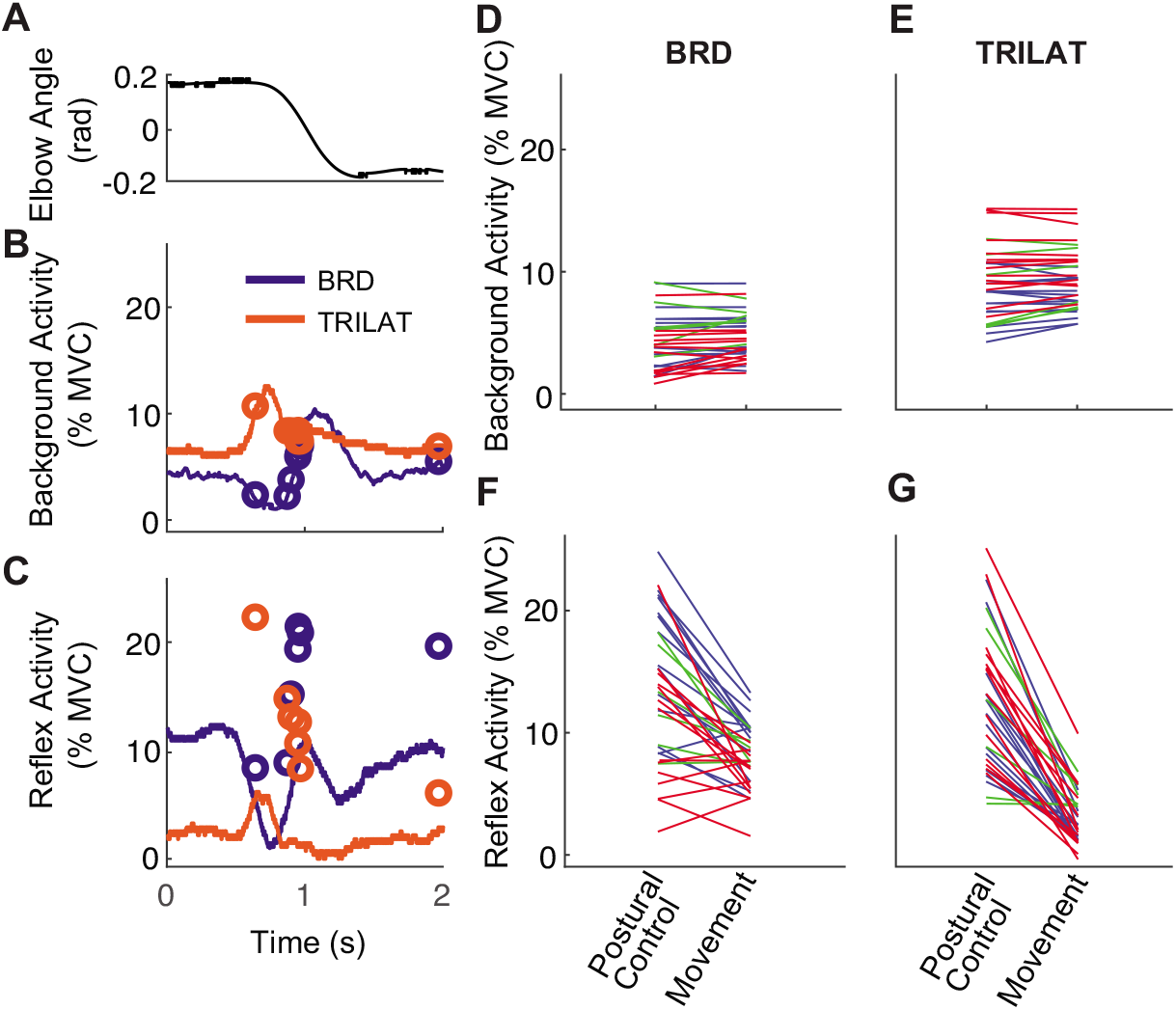
Differences in reflex amplitude for matched levels of background activity across tasks (postural control and movement) for the participants that completed both tasks (N = 4). **A –** Elbow angle from a representative subject during the extension phase of movement. **B** – curves show the background activity corresponding to the movement in **A**. Round markers illustrate the background activity recorded in 7 postural trials, designed to match the background activity during movement. Background activity was matched within 2 %MVC for each subject, resulting in a total of 34 paired comparisons incorporating both flexion and extension. **C** - Reflex activity corresponding to the conditions illustrated in **B**. **D,E –** Paired comparisons of the background activity during postural control and movement for the BRD and TRILAT, respectively. **F,G –** paired comparisons for the reflex amplitudes during postural control and movement.

The decrease in stretch reflex amplitudes during movement was not due solely to increased activity in the antagonist muscles. This was assessed by comparing the sensitivity of stretch reflexes in the BRD and TRILAT to the background activities in both muscles. The sensitivity of the BRD reflex to its own background activity increased by more than 3x to 2.03 ± 0.18 (t_3.3_ = 11.3, p < 0.001) during postural control compared to movement, while the magnitude of the sensitivity to its antagonist (TRILAT) background decreased by 0.44 ± 0.17 (t_2.0_ = 2.66, p = 0.12), to become insignificantly different than zero (t_1.6_ = −0.91, p = 0.48) (**Figure 7**). Similarly, the sensitivity of the TRILAT to its own background increased by more than 10x to 1.68 ± 0.17 (t_2.5_ = 9.80, p = 0.005). The magnitude of its sensitivity to the background of the antagonist increased by −0.62 ± 0.35 (t_3.0_ = −1.77, p = 0.17), but that difference did not reach statistical significance. These comparisons point to an increased sensitivity of the reflex to the background activity of the target muscle during postural control compared to movement. Additionally, we found no significant difference in the reflex sensitivity to the antagonist muscle’s background activity between the two tasks, though this can be attributed at least in part to the small numbers used in this control experiment.

**FIGURE 7.**
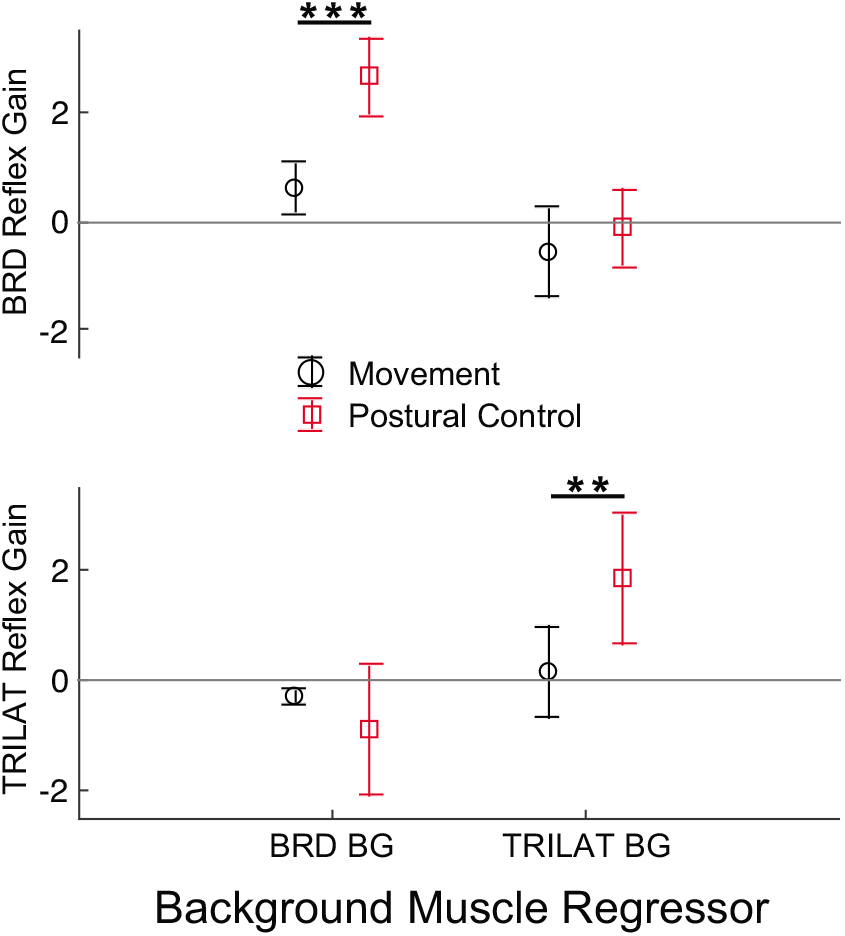
The effect of task (movement vs. postural control) on the gains of target muscle background and antagonist muscle background for a flexor and extensor reflex response for the participants that completed both tasks (N = 4). Error bars are 95% confidence intervals. ** signifies p < 0.01. *** signifies p < 0.001.

## Discussion

The goal of this study was to determine if the continuous modulation of short latency stretch reflex sensitivity during movement could be accounted for by the background activity in agonist and antagonist muscles relevant to movement control. We accomplished this by having subjects perform hundreds of elbow flexion and extension movements as a motor applied small random rotational perturbations to the elbow. We used surface EMG and a time-varying method to track both the background muscle activity and the perturbation-evoked reflex responses throughout the movement. We found that stretch reflexes during movement reflect the net volitional background activity in the agonist and antagonist muscles, and that models incorporating agonists and antagonists are significantly better than those considering only homonymous muscle gain-scaling. Increases in agonist muscle activity enhanced stretch reflex sensitivity whereas increases in antagonist activity suppressed reflex activity. Surprisingly, the magnitude of these effects was similar, suggesting a balance of control between agonists and antagonist that is very different that the dominance of sensitivity to agonist activity during postural tasks. This greater relative sensitivity to antagonist background activity during movement is due to a large decrease in sensitivity to homonymous muscle activity during movement rather than substantial changes in the influence of antagonist muscle activity. Together, our results provide a means to continually assess the coordination between volitional drive and reflex activity during movement and demonstrate that reflex sensitivity during movement reflects a balance of activity within agonist and antagonist muscles, unlike the predominant sensitivity to homonymous muscle activity in many postural tasks.

Several studies have demonstrated that the sensitivity of the short latency stretch reflex does not follow the target muscle’s background activity during movement, as would be expected with simple gain-scaling (Bawa, et al., 1999; Dufresne et al., 1980; Johnson M. T. et al., 1993; Mortimer et al., 1981; Soechting, et al., 1981). While these studies have related this modulation to the movement’s kinematic variables none looked at its relation to the net activity of the agonist and antagonist muscles controlling movement, which is best done when continuous estimates of reflex sensitivity are available. Dufresne, et al. (1980) used a time-varying method to obtain continuous measures of the reflex gain that are similar to ours, but did not assess the relationship between background muscle activity and changes in reflex sensitivity. Bawa, et al. (1999) assessed reflex modulation during movement with a velocity similar to ours. They also observed suppression of the stretch reflex during movement, they only measured reflex activity in time points of approximately every 250 ms. This time resolution is not sufficient to relate reflexes to antagonist background activity, for which an individual burst of activity may be as short as 150 ms during ballistic movements. In contrast, our paradigm of applying continuous perturbations and recording EMG at 2.5 kHz is sufficient to capture bursts of muscle activity associated with reflexes throughout the entire movement. To our knowledge, this is the first study investigating short latency stretch reflex amplitude with respect to antagonist muscle activity with sufficient time resolution to observe and model modulation during ballistic movements.

The reflex modulation we observed reflected a balance in the sensitivity of stretch reflexes to activity in agonist and antagonist muscles. This finding is similar to the balance of activity previously reported in muscle spindle responses to stretch (Dimitriou, 2014), suggesting that our results are at least in part due to control of spindle sensitivity. Such control would need to extend beyond *α-γ* coactivation or *β* innervation to include reciprocal inhibition of spindle firing (Sears, 1964). This reciprocal control, expressed at the level of whole muscle reflexes, represents a form of push-pull control (Johnson Michael D. et al., 2012) that could have a profound effect on the ability to regulate stretch reflexes during movement. Push-pull control would increase the range of achievable gains and rate of reflex modulation when antagonistic muscles are activated reciprocally, and could provide a mechanism for decoupling increases in reflex sensitivity from increases in agonist muscle activity through co-contraction. The extent of such use remains to be explored.

The balanced regulation of reflex activity we observed was due primarily to a decrease in gain of the homonymous stretch reflex compared to postural control. Several others have demonstrated a reduced sensitivity of the stretch reflex during movement (Bawa, et al., 1999; Ludvig, et al., 2017; Nakazawa, et al., 1997). Similarly, the H-reflex is also suppressed during movement (Brooke et al., 1992; Capaday et al., 1986; McIlroy et al., 1992), an attribute often attributed to presynaptic inhibition of the afferent pathways mediated through both descending drive (Capaday et al., 1987) and afferent feedback (Macefield, et al., 2018; Nielsen et al., 1995). Though we only conducted a limited comparison to postural control, our results suggest that presynaptic inhibition or other mechanisms that suppress stretch reflex gain scaling during movement are specific to homonymous stretch reflexes and have little effect on the sensitivity of reflexes to activity in the antagonist muscles.

### Limitations

While our methodology allowed for continuous estimates of stretch reflex activity during ballistic movements, it relied on the use of continuous perturbations that can suppress reflex activity (Pope et al., 2015). We limited the suppressive effects of continuous perturbations by using low mean absolute velocities (< 0.75 radians/s) that are outside the range traditionally associated with reflex suppression (Stein R. B. et al., 1995). The coefficients we estimated between background and reflex activity were also remarkably similar to the coefficients estimated in (Dimitriou, 2014) study on spindle afferents, suggesting that suppression of the stretch reflex was not a major factor in our experiments but it is not one that we can rule out completely.

### Conclusions

In conclusion, we developed an approach to characterize the continuous coordination between volitional drive to muscles the sensitivity of the stretch reflex. We found that the continuous changes in reflex sensitivity during movement were best explained by the balance of activity in the agonists and antagonists controlling movement. This balance was very different than what we observed in a postural task at matched levels of muscle activity, in which reflex sensitivity was due predominantly to the background activity in the homonymous muscle. These results demonstrate a push-pull configuration of stretch reflex control during movement, which may serve to heighten control and flexibility of stretch reflexes during movement, albeit at lower overall gains than during posture control. The methodology and results we have developed provide a framework for assessing the coordination of reflexive and volitional control across different tasks and pathological conditions.

